# A proteo-transcriptomic investigation of toxin evolution in planarians and their role in flatworm terrestrialization

**DOI:** 10.1101/2025.01.09.632098

**Authors:** Raquel García-Vernet, Lisandra Benítez-Álvarez, Javier Palma-Guerrero, Cristina Chiva, Klara Eleftheriadi, Iñaki Rojo, Eduard Sabidó, Rosa Fernández

## Abstract

The transition from aquatic to terrestrial environments represents a major evolutionary transition in animals, requiring significant adaptations in physiology and defense mechanisms to the challenges presented by the harsh terrestrial environment. Platyhelminthes, which include both aquatic and terrestrial species with a single terrestrialization event in the family Geoplaniidae, serve as excellent model organisms for studying the evolutionary adaptations required for terrestrialization. This study investigates the evolutionary dynamics of toxin orthologous groups (as a proxy to gene families) in aquatic and terrestrial flatworms, together with mucus composition, focusing on their role in terrestrialization from a molecular ecology perspective. Using a proteo-transcriptomic approach, we predicted and identified a broader toxin gene repertoire in terrestrial flatworms compared to freshwater ones. Although most toxins in flatworms arose before terrestrial planarian diversification—gaining a novel evolutionary origin at the Tricladida and Continenticola nodes—the mucus protein repertoire appears to have a far older evolutionary origin in both species. Moreover, distinct orthologous groups underpin the toxin gene repertoire and mucus composition in each lineage, highlighting the contrasting evolutionary trajectories of these two functional components. While toxin families in both aquatic and terrestrial flatworms revealed overall common functions, including cytokine modulation and ion channel regulation, terrestrial flatworms exhibited specific expansions of lectin-like proteins and pro-inflammatory responses, highlighting their potential key role to respond to land-based threats. This study provides new insights into the differential evolutionary trajectories of toxin and mucus proteins in planarians, offering a deeper understanding of the genetic innovations that facilitated flatworm terrestrialization.

## Introduction

The transition from aquatic to terrestrial environments marks one of the most significant events in evolutionary history, occurring independently across multiple animal phyla. This transition demanded profound physiological adaptations, including changes in locomotion, respiration, reproduction, predation, and defense against novel pathogens (Little, 1983). Genomic changes by gene gain, duplication and loss have accompanied these evolutionary shifts, driving the diversification of terrestrial lineages across various animal groups (Aristide & Fernández, 2023; Meyer et al., 2021). Some orthologous groups (as a proxy to gene families) that expanded or contracted in terrestrial species are likely directly related to overcoming the challenges posed by this transition, as observed for instance in aquaporins (Martínez-Redondo et al., 2023), directly involved in osmoregulation, and more expanded overall in freshwater and terrestrial lineages compared to marine ones.

A key challenge related to terrestrialization is the exposure to new pathogens, new predators and new prey. Toxins, typically peptides or proteins with harmful effects on other organisms, are key molecular players in predation and defense. These compounds have independently evolved from non-toxin proteins in at least eight different phyla (Casewell et al., 2013; Schendel et al., 2019). Despite their diversity, toxin molecular functions and domains often exhibit convergent evolution across lineages (Casewell et al., 2013; Fry et al., 2009; Zancolli et al., 2022). While venomous vertebrates and certain arthropod groups have traditionally received more research focus, recent advances in venomics are increasingly uncovering toxins in under-studied invertebrates (Jenner et al., 2019; Verdes et al., 2018, 2022). Notably, invertebrates, which lack adaptive immune responses (Kloc et al., 2024; Yacoub et al., 2020), also use toxins as a defense against pathogens, making them promising sources of antimicrobial peptides (Yacoub et al., 2020).

Despite this growing interest, the mechanisms driving toxin evolution across invertebrate lineages, particularly in response to terrestrialization, remain poorly understood. One promising group for studying these processes is the Platyhelminthes (flatworms). Terrestrialization within this phylum is confined to the monophyletic clade Geoplaniidae, part of the Tricladida order, which also includes freshwater and marine species (Riutort et al., 2012). The transition to land in planarians was accompanied by notable gene gains at the Tricladida node, which facilitated adaptation to terrestrial habitats (Benítez-Álvarez et al. 2025a). These gene repertoire changes likely played critical roles in addressing the new environmental challenges, including potentially toxin evolution.

Free-living flatworms are predators, and although all species utilize a muscular protrusible pharynx for feeding (Giribet & Edgecombe, 2020), hunting strategies vary between taxa. For example, while freshwater species like those in the Planariidae and Dugesiidae families often feed on weakened or immobile prey (Vila-Farré & C Rink, 2018), terrestrial species, such as those in the Geoplanidae, are active predators. These terrestrial flatworms use physical force and adhesive mucus to capture prey (Cuevas-Caballé et al., 2019; Sluys, 1999). In these species, mucus plays a key role in immobilizing prey, and it has been suggested to have neurotoxic properties (Thielicke and Sluys 2019; Sluys 2019; Cardoso et al. 2023), though the mechanisms remain unclear.

Planarians lack specialized venom glands, complicating the study of their toxins. However, toxins have been identified across the Platyhelminthes phylum, with most studies focusing on parasitic species. Toxins have also been reported in free-living flatworms, including tetrodotoxin (TTX) in some terrestrial species of the genus *Bipalium* (Stokes et al., 2014). In these species, TTX is concentrated in the head and eggs, similar to other marine flatworms (Miyazawa et al., 1987), suggesting region-specific toxin expression, particularly in body parts involved in hunting, such as the pharynx or secreted mucus, and offspring protection.

Although toxins may be localized in specific regions, expression patterns likely vary across species, and the absence of clear venom delivery systems or specialized venom-producing tissues makes toxin characterization challenging. Traditional methods like BLAST are insufficient for predicting protein toxicity (Sharma et al., 2022). However, recent advances in machine learning and deep learning provide more reliable predictions (Cole & Brewer, 2019; Sharma et al., 2022; Vishnoi et al., 2020), offering new avenues for toxin discovery.

In this study, we aim to characterize the toxin repertoire of a freshwater (*Schmidtea mediterranea*) and a terrestrial planarian (*Obama nungara*) using a proteo-transcriptomic approach, with the goal of further understanding how toxin diversity has evolved in response to different ecological pressures, particularly the transition from aquatic to terrestrial habitats. We employed two complementary methods: (i) *in silico* prediction of putative toxins and (ii) characterization of mucus secretions, focusing on identifying proteins with potential toxin activity. Additionally, we explored the evolution of putative toxins and mucus proteins across the Platyhelminthes phylum to understand their evolutionary origins and their roles in terrestrial adaptation, investigating how these proteins may have contributed to novel predatory strategies, defense mechanisms, and resistance to pathogens in land-dwelling species. This comprehensive approach sheds light on the molecular innovations that have accompanied the conquest of land within this animal lineage.

## Material and Methods

### RNA and protein extraction

Specimens of *Obama nungara* were collected in Sopelana, Vizcaya, while specimens of the freshwater planarian *Schmidtea mediterranea* were obtained from a lab culture established from specimens from Montjuic (Barcelona). All animals were maintained in starvation before sample processing (2-3 days for the *O. nungara* specimens; 7 days for the *S. mediterranea* specimens).

Each specimen was divided into three body parts (assigned as head, body and pharynx) by fixing first the specimens with RNAlater, dissecting with a scalpel, and freezing immediately with liquid nitrogen. Samples were stored at -80°C until the extraction. For *O. nungara*, a fragment from each body part was used per replicate for the RNA and protein extraction, while for *S. mediterranea* we combined four fragments from four different individuals per replicate. Both RNA and proteins were extracted by using TRIzol® reagent (Invitrogen, USA). The frozen samples were homogenized by adding 400μl of TRIzol® and using an electrical pestle until homogenous. The RNA was extracted by following the manufacturer’s instructions, while for proteins the TRIzol® protocol was modified (following (Simões et al., 2013)), and the final protein pellets were resuspended in 25μl of resuspension buffer (1%SDS and 8M Urea in Tris-HCl 1M, pH 8).

The mucus proteins were collected by submerging one specimen of *O. nungara* or *S. mediterranea* in 5% N-Acetyl Cysteine (NAC) in PBS buffer and incubating for 8 minutes, followed by the collection of the supernatant and overnight precipitation with 4 volumes of acetone at -20C, centrifugation at 20,000g for 20 minutes and resuspension in 50μl for *O. nungara* or 30μl for *S. mediterranea* of resuspension Buffer. For each species, we sequenced 3 replicates of each body region, including mucus.

The protein profiles of all the protein extracts were examined in a SDS page before preparing the sample for proteomic sequencing to make sure that there wasn’t any overrepresented protein masking other proteins. 4–20% Mini-PROTEAN® TGX™ Precast Protein Gels (Bio-Rad, Hercules, CA, US) were used with 10x Tris/Glycine/SDS as a running buffer. PageRuler™ Unstained Protein Ladder (Thermo Scientific, San Jose, CA, USA) was also loaded in the SDS page to serve as reference.

### De novo transcriptome assembly and differential expression analysis

The concentration of all samples was determined using the Qubit RNA BR Assay kit (Thermo Fisher Scientific). RNA libraries underwent polyA enrichment for library preparation and were sequenced on a NovaSeq 6000 platform (Illumina, 2 × 150 bp) to achieve a coverage of 6Gb.

Transcriptomes were assembled following the pipeline detailed in MATEdb (Fernández et al. 2022). In brief, adapters were removed with Trimmomatic 0.38 (Bolger et al., 2014), and the de novo assembly for each different species was performed using Trinity 2.11 (Haas et al., 2013). We used Transdecoder 5.5 (https://github.com/TransDecoder/TransDecoder) to perform the prediction of the coding regions, and we removed all non-metazoan sequences identified through Diamond 2.0.8 (Buchfink et al., 2015) and filtered with Blobtools (Challis et al., 2020). From these filtered predicted proteomes we also extracted the longest isoforms.

The differential gene expression analyses across different body parts were carried out following the Trinity pipeline using edgeR (Challis et al., 2020; Haas et al., 2013). Detailed parameters and scripts are provided in the Github repository.

### Sample digestion and proteomics sequencing

Samples (10 µg) were reduced with dithiothreitol (100 mM, 37 °C, 60 min) and alkylated in the dark with iodoacetamide (5 µmol, 25 °C, 20 min). The resulting protein extract was washed with 2M urea with 100 mM Tris-HCl and with 50 mM ammonium bicarbonate for subsequent digestion with endoproteinase LysC (1:10 w:w, 37°C, o/n, Wako, cat # 129-02541) and trypsin (1:10 w:w, 37°C, 8h, Promega cat # V5113) following Wiśniewski et al. FASP procedure. After digestion, the peptide mix was acidified with formic acid and desalted with a MicroSpin C18 column (The Nest Group, Inc) prior to LC-MS/MS analysis.

Samples were analyzed using a Orbitrap Fusion Lumos mass spectrometer (Thermo Fisher Scientific, San Jose, CA, USA) coupled to an EASY-nLC 1200 (Thermo Fisher Scientific (Proxeon), Odense, Denmark). Peptides were loaded directly onto the analytical column and separated by reversed-phase chromatography using a 50-cm column with an inner diameter of 75 μm, packed with 2 μm C18 particles (Thermo Fisher Scientific, cat # ES903).

Chromatographic gradients started at 95% buffer A and 5% buffer B with a flow rate of 300 nl/min and gradually increasing to 25% buffer B in 79 min and then to 40% buffer B in 11 min. After each analysis, the column was washed for 10 min with 100% buffer B. Buffer A: 0.1% formic acid in water. Buffer B: 0.1% formic acid in 80% acetonitrile.

The mass spectrometer was operated in positive ionization mode with nanospray voltage set at 2.4 kV and source temperature at 305°C. The acquisition was performed in data-dependent acquisition (DDA) mode and full MS scans with 1 microscans at resolution of 120,000 were used over a mass range of m/z 350-1400 with detection in the Orbitrap mass analyzer. Auto gain control (AGC) was set to ‘standard’ and injection time to ‘auto’. In each cycle of data-dependent acquisition analysis, following each survey scan, the most intense ions above a threshold ion count of 10000 were selected for fragmentation. The number of selected precursor ions for fragmentation was determined by the “Top Speed” acquisition algorithm and a dynamic exclusion of 60 seconds. Fragment ion spectra were produced via high-energy collision dissociation (HCD) at normalized collision energy of 28% and they were acquired in the ion trap mass analyzer. AGC was set to 2E4, and an isolation window of 0.7 m/z and a maximum injection time of 12 ms were used. Digested bovine serum albumin (New england biolabs cat # P8108S) was analyzed between each sample to avoid sample carryover and to assure stability of the instrument and QCloud (Chiva et al., 2018; Olivella et al., 2021) has been used to control instrument longitudinal performance during the project.

### Proteomics data analysis

Acquired spectra were analyzed using the Proteome Discoverer software suite (v2.5, Thermo Fisher Scientific) and the Mascot search engine (v2.6, Matrix Science). The data were searched against the newly-generated *de novo* assembled transcriptomes of *O. nungara* and *S. mediterranea* (see above) plus a list of common contaminants and all the corresponding decoy entries. For peptide identification a precursor ion mass tolerance of 7 ppm was used for MS1 level, trypsin was chosen as enzyme, and up to three missed cleavages were allowed. The fragment ion mass tolerance was set to 0.5 Da for MS2 spectra. Oxidation of methionine and N-terminal protein acetylation were used as variable modifications whereas carbamidomethylation on cysteines was set as a fixed modification. False discovery rate (FDR) in peptide identification was set to a maximum of 1%.

Peptide quantification data were retrieved from the “Precursor Ions Quantifier” node in Proteome Discoverer (v2.5) using a mass tolerance of 2 ppm for the peptide extracted ion current (XIC). The obtained values were then used to calculate the Top3 value for each protein, which is the average of the three most abundant peptides for each protein.

The final list of identified proteins of each species was constructed by merging the results of the 12 samples (3 replicates for each body part, including the mucus fraction), but only proteins identified with high confidence levels and designated as the lead protein within a protein group (master protein) were considered. A protein group was defined as a collection of proteins containing at least one unique peptide. We finally explored the relationships among sample replicates through a Principal Component Analysis (PCA) using as input the normalized abundances of all the master proteins. All statistical analyses were performed using Proteome Discoverer software suite.

For specifically characterizing the mucus of each species, we selected a subset of proteins that were differentially abundant in the mucus, in comparison to the body samples, that were used as the background signal. In the protein subset we included (i) proteins that were differentially expressed between the mucus and the body background signal, exhibiting a significant adjusted p-value and higher abundance in the mucus, and (ii) proteins that were identified in at least 2 out of 3 samples of the mucus, and were not present in non of the remaining samples (i.e., proteins that were detected as unique in the mucus fraction).

### Prediction of toxins and functional annotation

The predicted proteomes of each species were used to predict candidate toxins. The candidate toxins were predicted by using two different softwares: Toxify, which follows a deep learning approach (Cole & Brewer, 2019) and Toxinpred2, which follows an hybrid method that combines BLAST-based similarity, motif search and prediction models (Sharma et al., 2022), with a threshold of 0.6 and 0.95 respectively. Candidate toxins predicted by both softwares were merged to generate a list, which was used to detect the secreted toxins by using SignalP 6.0. To reduce the amount of false positives during toxin prediction, we compared the list of putative secreted toxins with the proteins identified in the proteomic analyses, and built a final list including only those toxins that were also detected in the proteomic analyses.

The putative toxins and the subset of mucus proteins were annotated through BLAST, using as reference database the non-redundant protein sequences (nr). Since many putative toxins yielded no hits, we further explored putative protein function with FANTASIA (Martínez-Redondo et al. 2024), which leverages natural language models to infer Gene Ontology (GO) terms. Finally, we explored which GO terms were enriched in the mucus and toxins subsets against the proteome of each species inferred through proteomics using TopGO (Alexa & Rahnenfuhrer 2024) and REVIGO 1.8.1 (Supek et al., 2011), with the Gene Ontology database dated from january 2024. Graphic representations were done using RawGraph (Mauri et al. 2017) and (R version 4.2.2)

### Toxins and mucus evolution across the phylum Platyhelminthes

We used the dataset and methods described in Benitez-Álvarez et al. 2025a and Benítez-Álvarez 2025b. Briefly, we combined the longest isoforms from the de novo transcriptomes of *O. nungara* and *S. mediterranea* (this study), with data from 31 other platyhelminthes species (Benítez-Álvarez et al. 2025b). We also included 4 outgroups belonging to the phyla Mollusca, Nemertea, Annelida, and Gastrotricha (Benítez-Álvarez et al. 2025b), obtaining a final dataset of 37 species. We inferred the Hierarchical Orthologous Groups (HOGs) with OMA v2.6 (Altenhoff et al., 2013), identifying the origin of HOGs at each diversification event in the phylogeny through pyHam (Train et al., 2019).

For all the HOGs containing genes identified as toxins only for one species, but also sequences from the other species we explored potential shifts in the selection pressure using Pelican (Duchemin et al., 2023). We coded the aquatic lineages as background and terrestrial ones as foreground characters. A p-value at gene level was estimated using using the Gene-wise Truncated Fisher’s method considering the best k=10 p-values in each alignment (https://gitlab.in2p3.fr/phoogle/pelican/-/wikis/Gene-level-predictions). Additionally, we applied the adjustment method Benjamini & Hochberg (BH) (Benjamini & Hochberg, 1995) based on the false discovery rate to obtain an adjusted p-value.

Finally, we used the IHam tool (Train et al., 2019) to visually inspect the clustering of gene members across all HOGs that included toxins and mucus sequences. From this inspection, we identified the HOGs that showed expansions or contractions in the terrestrial lineage. Detailed methods and data from the species included in these analyses can be found in the Github repository and Benitez-Álvarez et al. 2025a, Benítez-Álvarez et al. 2025b.

## Results

### Toxin prediction and proteomic identification

For each species, we constructed a high-quality reference transcriptome and predicted its proteome by merging the 9 RNA libraries (see Github repository). These libraries corresponded to different body regions, including the head, pharynx, and the rest of the body, for both *O. nungara* and *S. mediterranea*.

Based on the predicted proteome, we identified putative toxins using Toxify (Cole & Brewer, 2019) and ToxinPred2 (Sharma et al., 2022). ToxinPred2 predicted a greater number of toxins than Toxify. Regardless of the software used, we detected a higher number of toxins in *O. nungara* than in *S. mediterranea* (Figure 1A, 1B, Github repository).

**Figure 1.**
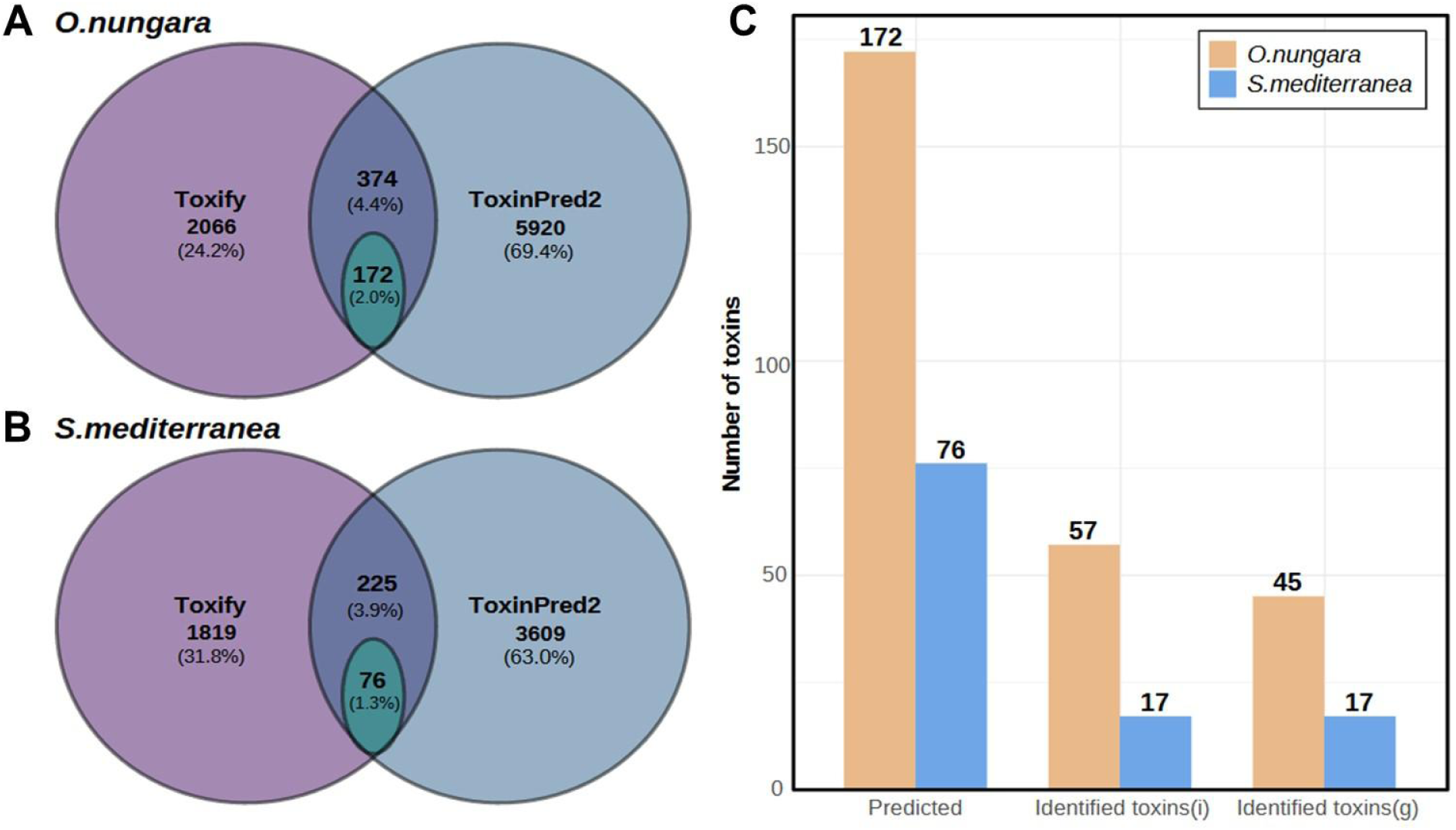
Toxin predictions and identifications. The Venn diagram for (**1A**) *Obama nungara* and (**1B**) *Schmidtea mediterranea*, showing the number of toxins predicted by Toxify (purple) and ToxinPred2 (blue). Green circles represent proteins predicted to be secreted from the shared list of toxins identified by both software tools. **1C.** Barplot showing the proteins predicted and identified for *Obama nungara* (brown) and *Schmidtea mediterranea* (blue). “Identified toxins (i)” refers to the total amount of toxins identified in the proteomic analyses, while "Identified toxins (g)" indicates the number of distinct genes corresponding to these toxins.

For both species, we performed a protein extraction from the previously mentioned body regions as well as the mucus, and sequenced them through a bottom-up proteomics approach. We unambiguously identified 3584 proteins in *S. mediterranea* and 5302 proteins in *O. nungara* (Github repository). To reduce false positives in our toxins predictions, we compared our final toxin candidates with the proteins identified in the proteomic analyses. Again, the number of toxins found was higher in *O. nungara* than in *S. mediterranea* (Figure 1C). Most of the toxins that were identified in the proteomic data corresponded not only to different isoforms, but also to different genes.

### Toxin characterization

BLAST results revealed that, although functional annotations were similar in both species, the most frequent toxins differed between the aquatic and terrestrial planarians (Figure 2A, Table S1). In *O. nungara*, 12 out of 57 toxins were annotated as lectins or scavenger receptors, whereas this function was not found in any of the toxins predicted in *S. mediterranea*. Two functions were also exclusively found in *S. mediterranea*, which included one conotoxin and one follistatin. On the other hand, both species contained toxins annotated as phospholipases, insulin-like proteins, granulins, and venom allergens. Finally, for a considerable proportion of toxins, we obtained no match or found the function to be unknown. In addition, some annotations were either too general or unclear.

**Figure 2.**
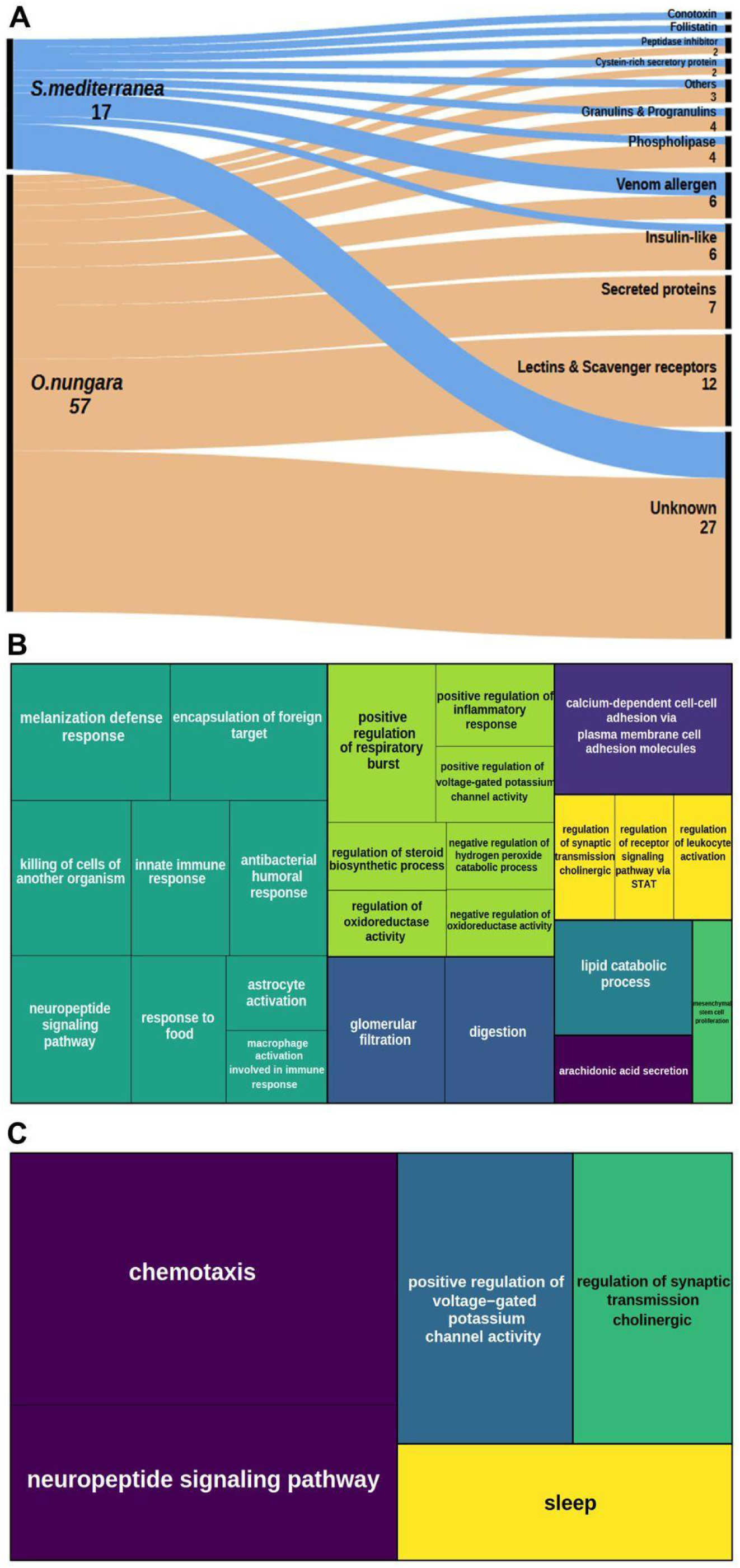
Functional annotation of all the putative toxins identified in the proteomic results. **2A.** Alluvial plot showing the summarized BLASTp annotation results, with *O. nungara* in brown and *S. mediterranea* in blue. Numbers indicate the amount of toxins included in a specific category. **2B, 2C.** Results of the enrichment analyses for the Biological process category in *O. nungara* (B) and *S. mediterranea* (C).

Enrichments performed with FANTASIA annotations showed that the amount of enriched categories was substantially higher in *O. nungara* than in *S. mediterranea*, likely reflecting the greater amount of toxins predicted in the terrestrial planarian. In general, most of the terms that were enriched in *S. mediterranea* also were found enriched in *O. nungara*.

Although in the biological process category three out of five categories were common between both species, *O. nungara* exhibited a notable number of categories related to the immune system, defense mechanisms, and hunting, such as “killing cells of another organism”, “antibacterial humoral response”, or “melanization defense response”. We also identified categories related to respiration and oxidative stress (indicated by pale green squares). In *S. mediterranea* we found an additional category related with the nervous system (“sleep”), and also “chemotaxis” (Figure 2B,C).

Regarding the molecular function (Figure S1), both species shared two common enriched categories: "cytokine activity" and "ion channel regulation activity", in concordance with the findings from the biological process enrichments. *S. mediterranea* had only one additional enriched GO term, "toxin activity". In contrast, in *O. nungara* we detected up to 10 more enriched GO terms, including among others several binding-related categories, such as “galactose binding” or “antigen binding”, as well as immune system-related categories.

To fully characterize the function and expression of these putative toxins, we examined whether there was an increased expression across the various body parts analyzed in both transcriptomic (head and pharynx) and proteomic data (head, pharynx and mucus) when compared against the body background signal. We found only one toxin that was significantly differentially expressed in the pharynx of *S. mediterranea* in the transcriptome (identified as “SMED3_DN4834_c0_g1”). This same protein showed higher abundance also at protein level, although the difference was not significant (Table S2). BLAST annotations did not retrieve any hits for this protein, but FANTASIA annotations indicated a potential relationship with the neuropeptide signaling pathway and some kind of calcium binding activity (Github repository).

At proteomic level, we did not find any putative toxin which was significatively more abundant in the regions analyzed compared with the body background signal. However, for both species, we found some toxins showing relatively higher concentrations in the head, mucus or pharynx, reflected in the abundance ratio of each region compared to the body background signal (Table S2, Table S3). Nonetheless, the normalized abundance values across replicates exhibited a considerable variability, and in some cases there was also a lack of quantification in samples from specific body regions. Despite the observed trends, with our results we cannot confirm nor discard differential expression of the putative toxins across the body of *S. mediterranea* and *O. nungara*, and further studies will be needed to determine if true differential expression exists.

### Mucus protein characterization

SDS page gel showed that the mucus fraction presented a different protein composition pattern in comparison with the other body fractions (Figure S2), and PCAs revealed a similar result (Figure S3). To go further in this, we created mucus protein subsets for each species containing (i) proteins that were significantly differentially expressed in mucus, and (ii) proteins that were only found in mucus samples. While our strict criteria aim to exclude proteins that are also abundant in the body of these animals - and therefore potential contaminants produced during the extraction of the mucus fraction - our subsets may not capture the full mucus composition. Instead, our subsets primarily include the most abundant and specific proteins in *O. nungara* and *S. mediterranea* mucus.

Enrichment analyses of both species retrieved terms related to adhesion processes, as well as terms related with development, growth, regeneration and immune system (Figure 3A, B). In both species, we annotated through blast proteins that were related with cell structure (such as actin, myosin and other related proteins), adhesion (mucins, von Willebrand factor), immune system and defense (lectins, proteases) and proteins related with metabolism and nucleic acids (such as ribosomal proteins). Full results are detailed in the Github repository and Figure S4.

**Figure 3.**
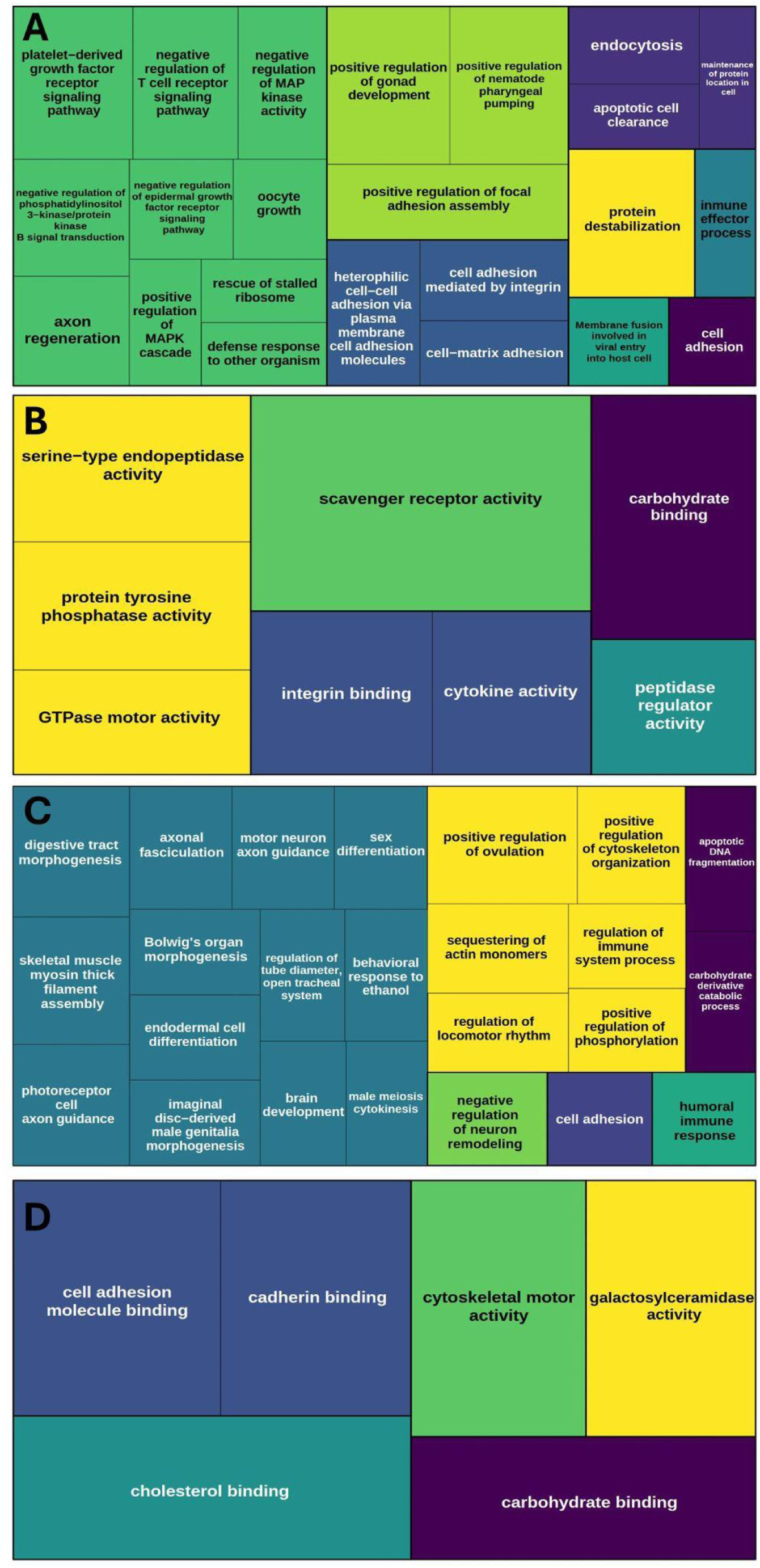
Results of the enrichment analyses of mucus for *O. nungara* in the Biological process (A) and Molecular function (B) and *S. mediterranea* in the Biological process (C) and Molecular function (D).

In addition to these conserved general patterns, in *O. nungara* the enrichments revealed GO terms that coincided with the most common toxin annotations, including ’scavenger receptor activity,’ ’carbohydrate binding,’ and ’cytokine activity’, as well as a higher amount of GOterms related with immune system and defense, such as “defense response to other organism”, “immune effector process” or “protein destabilization”, which could also be related with the lectins identified in the mucus fraction. On the other hand, enrichment analyses of *S. mediterranea* retrieved a higher number of GO terms that were related with morphogenesis and development (Figure 3).

### Evolutionary origin of toxins and mucus proteins

To better understand the role of mucus and toxins during the transition from aquatic to terrestrial environments, we explored the evolutionary origin of toxin- and mucus-protein orthologous groups across the platyhelminth phylogeny. The general results of gene assignment to hierarchical orthologous groups (HOGs) are outlined in Table 1.

**Table 1:** Number of genes of each species (*O. nungara* and *S. mediterranea*) and dataset (toxins and mucus) and its assignment to hierarchical orthologous groups (HOGs). Genes that were species-specific were not assigned to any HOG. Finally, the number of HOGs that were gained after the diversification of terrestrial planarians (*O. nungara* results) and freshwater planarians (*S. mediterranea* results) are detailed.

| Subset of genes | Species | Genes | Species-Specific | HOGs | Terrestrial / Freshwater unique |
| --- | --- | --- | --- | --- | --- |
| Toxins | *O. nungara* | 45 | 5 | 24 | 8 |
| Toxins | *S. mediterranea* | 17 | 0 | 14 | 1 |
| Mucus | *O. nungara* | 102 | 13 | 87 | 21 |
| Mucus | *S. mediterranea* | 97 | 6 | 87 | 6 |

A high proportion of HOGs including predicted toxins and mucus proteins originated prior to the diversification of terrestrial planarians, although mucus and toxin orthologous groups showed different patterns (Figure 4). A large proportion of toxin orthologous groups was gained in the branch leading to Tricladida (which includes marine, freshwater and terrestrial species) and Continenticola (Github repository). On the other hand, most mucus orthologous groups of *O. nungara* and *S. mediterranea* had an ancient origin, with approximately half of the total HOGs emerging before the diversification of platyhelminthes and therefore being shared with other animal phyla.

**Figure 4.**
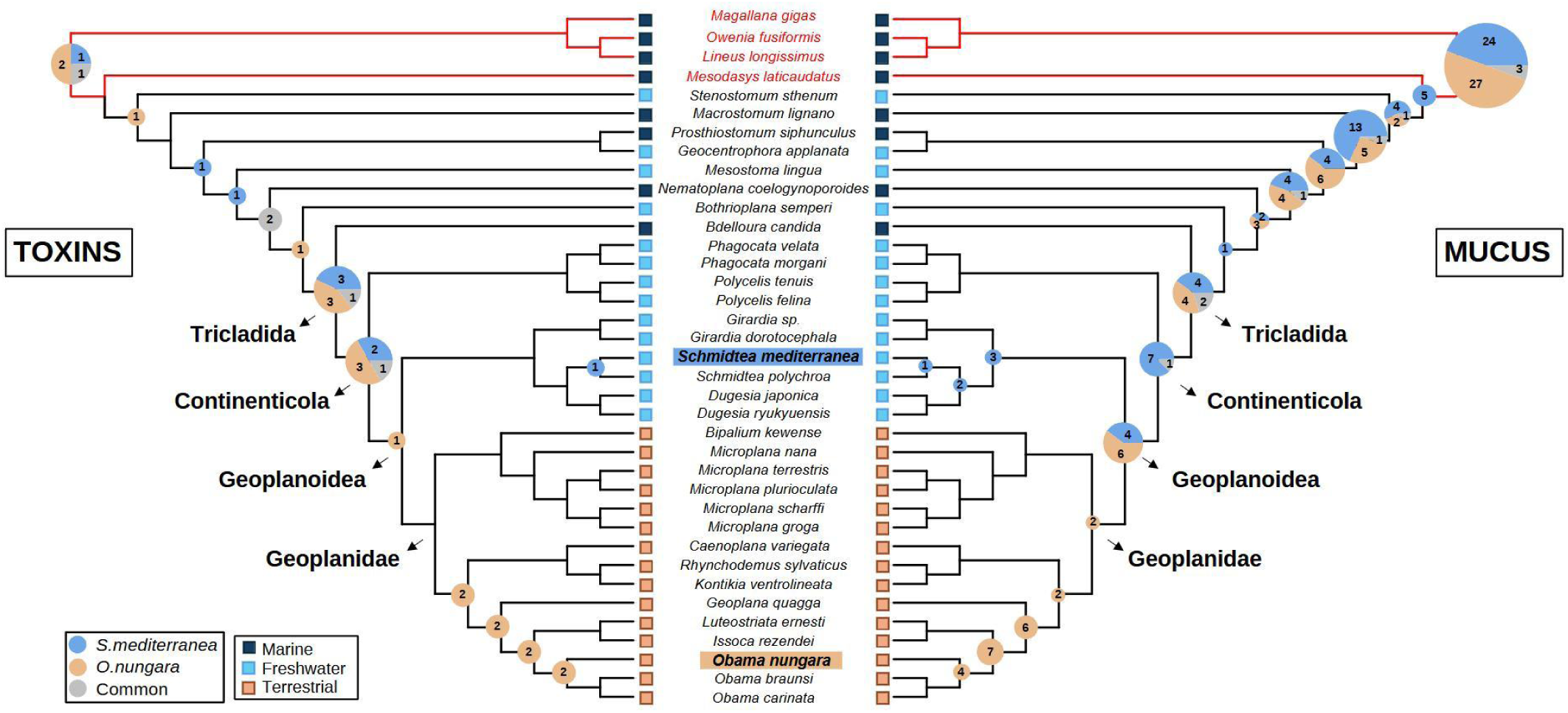
Evolutionary origins of toxins and mucus proteins across Platyhelminthes. Outgroups are highlighted in red, and the habitats of the different species are indicated by squares: dark blue for marine, light blue for freshwater, and brown for terrestrial. Gained HOGs are represented by circles at the nodes, with the colors and numbers inside indicating HOGs that contain sequences from *S. mediterranea* (blue), *O. nungara* (brown), or both species (grey).

**Figure 5.**
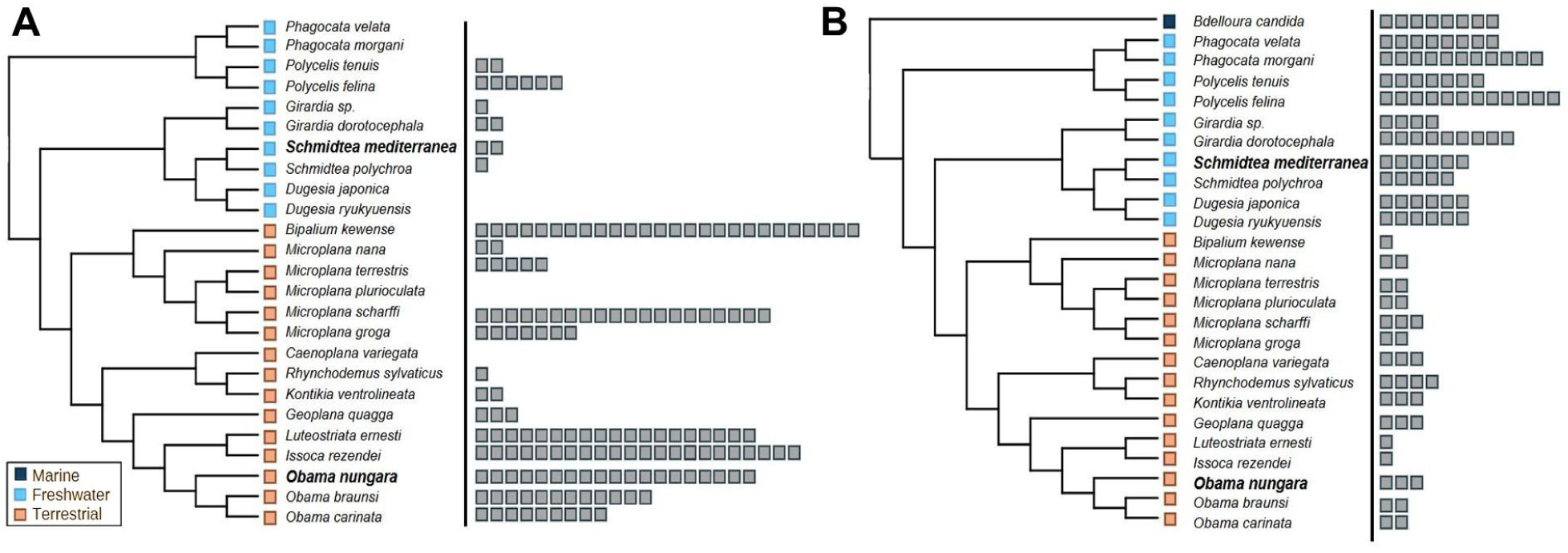
HOGs expanded or contracted in terrestrial flatworms. The habitats of the different species are represented by colored squares: dark blue for marine, light blue for freshwater, and brown for terrestrial. The number of gray squares aligned with each species name corresponds to the number of gene copies that the species has in the specified HOG. **5A**. HOG (HOG00036620_64) is expanded in terrestrial flatworms. **5B**. HOG (HOG00044669_65) is contracted in terrestrial flatworms.

Interestingly, despite the majority of the orthologous groups originated before the terrestrial planarians diversification, most of the HOGs were not shared between species, indicating that *O. nungara* and *S. mediterranea* may rely on different orthologous groups in their toxin and mucus composition. However, while mucus composition was assessed by direct identification in the selected subset of proteins, and therefore reflects the most abundant and specific proteins in the mucus of each species, the toxin identification was based on in silico predictions. In order to further clarify this, we conducted a deeper investigation into the sequences within the HOGs containing toxin genes to determine whether the HOGs with predicted toxins in one species (*O. nungara* or *S. mediterranea*) also included orthologous sequences from the other species. Five out of 33 HOGs contained toxins that were both predicted and identified in both species. However, of the remaining 28 HOGs, we identified 15 that included toxins predicted and identified in only one species (*O. nungara* or *S. mediterranea*), yet also contained sequences from the other species, which in most cases were not predicted as toxins by Toxify (see TableS4 for details). Results obtained with Pelican showed that 10 out of these 15 HOGs exhibited a significant shift in the selection at the amino acid level between aquatic and terrestrial planarians (Table S5).

We also detected two HOGs containing mucus genes that specifically originated in the ancestor of terrestrial planarians. The first HOG contained up to 10 *O. nungara* genes, including at least one identified in the mucus subset (“ONUN2_DN347_c0_g1”). GO terms inferred with FANTASIA were again associated with immune system response, retrieving some interesting terms as “defense response to bacterium” or “innate immune response” (Github repository). The other HOG gained in the ancestor of terrestrial planarians contained only one *O. nungara* gene. In this case, annotations included functions as “lipid binding” and also functions related with entry to the cell (Github repository).

Finally, we used pyHam to visualize the toxin and mucus HOGs and to identify potential expansions and contractions that could be associated with the adaptation of the planarians to the terrestrial environment (Github repository). We did not identify any mucus orthologous groups that presented expansions or contractions in terrestrial planarians. However, among the HOGs containing toxin genes, one showed an expansion and another a contraction in terrestrial planarians. The first orthologous group originated in the Continenticola node and was expanded in terrestrial planarians, with 19 gene copies in *O. nungara* and up to 26 copies in the case of the terrestrial flatworm *Bipalium kewense* (Figure 4A). At least 6 sequences of *O. nungara* were predicted as secreted toxins by the two software tools and identified in the proteomic results. Interestingly, and in concordance with previous results, all the BLAST hits of these sequences retrieved hits with lectins and scavenger receptors (Table S1).

The second HOG was gained in the branch leading to Tricladida and contracted in the terrestrial planarians (Figure 4B). Two out of the 6 sequences of *S. mediterranea* included in this HOG were predicted as secreted toxins, with one being also found in the proteomic results (Table S4). The BLAST annotation indicated that this toxin was classified as a venom allergen. Summarized results for all the proteins classified as toxins and mucus components, including the HOGs to which they belong and a general category for the BLAST hits can be found in Table S1 and S6.

## Discussion

Our study provides new insights into the composition and evolutionary trajectories of toxins and mucus proteins in flatworms and their potential role in flatworm terrestrialization. By comparing the terrestrial flatworm *O. nungara* and the freshwater species *S. mediterranea*, we aimed to elucidate how the adaptive gene repertoires differ between aquatic and terrestrial species, particularly in relation to toxins and mucus proteins.

### Evolutionary origin of the toxin repertoire in aquatic and terrestrial flatworms

We found that a substantial proportion of the predicted toxin families in both species arose before the diversification of the flatworm families examined here (Dugesiidae and Geoplanidae). This observation is consistent with recent findings indicating that many genes differentially expressed under abiotic stress conditions also originated prior to their diversification, specifically in the branches leading to Tricladida and Continenticola (Benítez-Álvarez et al. 2025a). Taken together, these results suggest that the genetic framework for toxin production was largely established earlier in flatworm evolution than previously thought, providing a reservoir of ancestral genes that could be co-opted or fine-tuned in subsequent lineages. This ancient gene repertoire may have played a pivotal role in enabling flatworms to adapt to diverse ecological niches, including both freshwater and terrestrial habitats, by conferring pre-existing molecular mechanisms for defense, prey capture, or other adaptive functions.

Despite a common evolutionary origin mostly in the branches leading to Tricladida and Continenticola, most of the toxin-containing HOGs were not shared between species. However, a substantial proportion of HOGs that included toxins predicted for only one species also contained sequences from the other. These results can be explained by two potential hypotheses: (i) although our protocol aimed to reduce the proportion of false positives by combining the results of two different prediction software tools, this approach may have increased the proportion of false negatives. Consequently, some sequences not predicted as toxins could indeed exhibit toxin activity; (ii) species may rely on distinct HOGs for their toxin repertoire, and genes belonging to the same HOG may have different functions depending on species-specific or environmental factors. In line with this, selection analyses revealed that approximately 66% of these HOGs exhibited a significant shift in selection between terrestrial and aquatic species, supporting the possibility that, at least in some cases, the latter hypothesis could be true. Hence, these findings suggest that differences in selection pressures may shape the specialized toxin repertoires of *O. nungara* and *S. mediterranea*, underscoring the complexity of molecular adaptations that emerge as flatworms occupy different environments. However, our data do not allow us to clearly distinguish between these two explanations, and further experimental studies will be required to definitively determine whether these proteins have indeed toxin activity.

### A larger toxin gene repertoire in terrestrial flatworms and its potential role in land colonization

We detected more putative toxin-related genes in *O. nungara* than in *S. mediterranea,* also reflected in the greater enrichment of significant categories in the terrestrial species. Specifically, the higher number of enriched GO terms related to immune response and inflammation in *O. nungara* suggests that these functions could have played a significant role in the adaptation to terrestrial life, where exposure to new pathogens, predators, and environmental stressors may have produced a more complex interaction between toxins and terrestrial ecosystems.

Although many toxin protein families were detected in both species, we observed some characteristic patterns which differed between the terrestrial and aquatic flatworms. For example, in *O. nungara*, at least 12 predicted toxins were identified as lectin-like proteins. Lectins are proteins that can bind to carbohydrates, and previous research suggests that these proteins may have a key role in the pathogen recognition and the immune system of invertebrates (Hanington et al., 2010; Pees et al., 2016) and have been also identified in venomous animals from other phyla including cnidarians, mollusks and nemertea (Delgado et al., 2022; Verdes et al., 2022; von Reumont et al., 2020).

The 12 lectins identified in *O. nungara* belonged to two different HOGs: the first originated within the terrestrial planarian lineage, while the second was gained in the branch leading to Continenticola and underwent a significant expansion in terrestrial species, (including *O. nungara*). This expansion aligns with previous observations in mollusks, where several C-type lectin families are expanded in terrestrial species and have been suggested to serve defensive roles in terrestrial environments (Aristide & Fernández, 2023) In addition, in planarians, several novel lectins have been also reported in freshwater flatworms (Gao et al., 2017; Shagin et al., 2002). One of these novel lectins, described in the freshwater planarian *Dugesia japonica*, has been associated with innate immune response against pathogens and with the first steps of regeneration and wound healing (Gao et al., 2017), further supporting its potential role in the adaptation to new environments and pathogens and its involvement in regeneration.

We also identified HOGs containing group 1 (secreted) Venom Allergen-like proteins (VAL), which are known to be widely distributed across metazoans, including flatworms (Chalmers et al., 2008). One of these families has an ancient origin and is shared across multiple phyla. The other two emerged at the branch leading to Tricladida. One of these HOGs experienced a contraction in terrestrial species, while the aquatic planarians, including *S. mediterranea*, showed a larger copy number. While the role of VAL proteins has been linked to parasitism and their function has been mainly studied in parasitic species (Chalmers & Hoffmann, 2012; Yoshino et al., 2014), the most expanded VAL orthologous group of this phylum has been precisely described in freshwater free-living flatworms (Chalmers & Hoffmann, 2012,Sipley 2019), in concordance with our results. Altogether, it seems that VAL proteins may have a relevant role in the biology of aquatic free-living planarians, with a potential shift in functional roles as flatworms adapted to terrestrial life.

### Functional convergence of toxins with different evolutionary origin in aquatic and terrestrial planarians

Beyond the functional differences between the toxin gene repertoire in aquatic and terrestrial flatworms discussed above, our analyses revealed that several functions identified in *O. nungara* and *S. mediterranea* were shared between both species, despite being encoded by different HOGs. Common enriched categories included cytokine activity, ion channel regulation and categories related with the neuromuscular system, suggesting that, as observed in other species and phyla, some flatworm toxins may trigger inflammatory responses, target ion channels and/or disrupt the nervous system (Brodie, 2009; Zhang, 2015).

In both species, we identified several toxins belonging to the CAP superfamily (Gibbs et al., 2008), including one conotoxin-like protein in *S. mediterranea* and Venom Allergen-like (VAL) proteins in both species, the latter of which have previously been described in different flatworm lineages (Chalmers & Hoffmann, 2012). CAP-domain proteins have been convergently recruited in venoms of many taxa, ranging from mammals to annelids or arthropods (Jenner et al., 2019; Verdes et al., 2018; von Reumont et al., 2020; Zhang et al., 2022). These proteins exhibit several biological functions, including disruption of ion channels, proinflammatory activity and the induction of allergic responses (Fry et al., 2009; Zhang et al., 2022)

We also identified phospholipases, which are frequently recruited in venoms of other taxa (Lynch, 2007; Undheim et al., 2014; Verdes et al., 2018) and induce multiple effects on the targets, including myotoxicity, neurotoxicity, inflammation and changes in the reactive oxygen species production (El-Benna et al., 2021; Fry et al., 2009). Finally, insulin-like proteins were also predicted and detected in both species. This protein group is also found in venoms across various taxa, including nemerteans (Verdes et al., 2022; von Reumont et al., 2020) and cone-snails (Guo et al., 2024), where it is used to facilitate prey capture.

### Ancient origins of mucus proteins encoded by distinct HOGs in aquatic and terrestrial flatworms

In contrast to toxin genes, the mucus protein repertoire appears to have a more ancient evolutionary origin, with many mucus-related HOGs also present across other animal phyla. Despite this shared ancestry, *O. nungara* and *S. mediterranea* seem to rely on distinct sets of HOGs for mucus composition, suggesting that each species draws upon different genetic resources to produce its most abundant mucus proteins.

For example, although we found proteins and GO terms related to immune defense in both species, the proportion of those in *O. nungara* mucus was higher, which could be due to an enhanced protective function of the mucus in terrestrial habitats. In both species, we found proteases and protease inhibitors (such as serpin proteins), including metalloproteases in the *S. mediterranea* mucus. Both proteases and proteases inhibitors have been repeatedly reported in the mucus of different species (Cerullo et al., 2023; Pales Espinosa et al., 2016), including *S. mediterranea (Bocchinfuso et al., 2012)*, and it has been suggested that these proteins contribute to the immunity. We also detected lectins in the mucus of both species, aligning with previous studies in planarians (Shagin et al., 2002; Zayas et al., 2010) and other invertebrates, where lectins have been suggested to contribute to the mucus adhesion and microbial protection (Liegertová et al., 2022; Smith et al., 2021).

Additionally, proteins involved in adhesion, such as mucins, spondin and fibrillin-like proteins, were present in the mucus of both species, underscoring their functional significance in aquatic and terrestrial habitats (Lang et al., 2007; Liegertová & Malý, 2023). Finally, in both species, we also detected proteins of intracellular origin, including those related with cell structure and cytoskeleton, such as actin or myosin, and ribosomal proteins. These results are highly consistent with previous mucus characterizations in species from other phyla (Lopes et al., 2024; Pales Espinosa et al., 2016; Schwaner et al., 2024) and have been also previously described in the *S. mediterranea* mucus (Bocchinfuso et al., 2012).

### Conclusions

Overall, we successfully predicted and identified putative translated toxins in *O. nungara* and *S. mediterranea,* and characterized the most abundant proteins in their mucus secretions. Our findings revealed both similarities and contrasting dynamics in the toxins and mucus proteins of these two flatworm species, which we discuss in the context of flatworm terrestrialization. We predicted a larger number of toxins in *O. nungara*, also reflected in a higher number of enriched GO terms related to immune response and inflammation. In both species, most orthologous groups encoding its mucus and toxin repertoire originated before the major split of terrestrial flatworms, showcasing how genetic elements that originated prior to the colonization of terrestrial environments were co-opted to deal with threats related to life on land. However, while most toxin-related HOGs were gained in the branches leading to Tricladida and Continenticola, a significant proportion of mucus-related HOGs seem to have a more ancient evolutionary origin. Although the toxin families in *O. nungara* and *S. mediterrane*a appear to share similar functions, terrestrial flatworms exhibit an expansion of a lectin orthologous group, suggesting that some lectin families may have a role in facilitating adaptation to land. These findings underscore the pivotal role of ancient genetic frameworks, supplemented by lineage-specific expansions like those of lectin families, in shaping the evolutionary success of flatworms on land. Altogether, these results highlight the importance of examining both ancient genomic elements and lineage-specific expansions to better understand how flatworms—and, by extension, other taxa—have successfully adapted to life on land.

## Data availability

Full link to the Github repository: https://github.com/MetazoaPhylogenomicsLab/Garcia_Vernet_et_al_2024_Flatworm_toxin_evol ution

## Acknowledgements

R.G-V was funded through a Margarita Salas grant, funded by the Spanish Ministry of Universities and the European Union Next Generation EU/PRTR. RF acknowledges support from the following sources of funding: the European Research Council (this project has received funding from the European Research Council (ERC) under the European Union’s Horizon 2020 research and innovation programme (grant agreement no. 948281), the OSCARS project (funding from the European Commission’s Horizon Europe Research and Innovation programme under grant agreement no. 101129751) and the Secretaria d’Universitats i Recerca del Departament d’Economia i Coneixement de la Generalitat de Catalunya (AGAUR 2021-SGR00420 and 2021-SGR01225). We acknowledge support of the Spanish Ministry of Science and Innovation through the Centro de Excelencia Severo Ochoa (CEX2020-001049-S grant funded by MCIN/AEI/10.13039/501100011033). The CRG/UPF Proteomics Unit is part of the Spanish Infrastructure for Omics Technologies (ICTS OmicsTech). We also thank Centro de Supercomputación de Galicia (CESGA) and the HPC Drago from the Centro Superior de Investigaciones Científicas for access to computer resources.

## Supplementary information

**Figure S1:** Results of the enrichment analyses for the Molecular function category in *O. nungara* and *S. mediterranea*.

**Figure S2:** SDS Page results for the different body fractions sampled in *O. nungara* and *S. mediterranea*. 10 micrograms of protein per lane. A. SDS Page coomassie stained. Lane 1, PageRuler™ Unstained Protein Ladder (ThermoScientific), Lanes 2 to 5 proteins from *O. nungara*. Head shown in lane 2, body in lane 3, pharynx in lane 4, and mucus in lane 5. Lanes 6 to 10 proteins from S. mediterranea. Head in lane 6, body in lane 7, pharynx in lane 8, lane 9 and 10 shows mucus extracted by 4 minutes or 8 minutes incubation in NAC respectively, lane 10. Note that the mucus fraction in lane 5 is from a failed attempt of collecting mucus directly by scrapping the individuals, so we repeated it with NAC as for *S. mediterranea* and it is presented in B. B. SDS Page of *O. nungara* proteins from mucus and body. Lane 1 shows the protein ladder. Lanes 2 and 3 show the mucus extracted by 4 minutes or 8 minutes incubation in NAC respectively. Lane 4 shows protein from the body.

**Figure S3:** PCAs showing the distinct proteomic compositions of the body fractions (body, head, mucus, and pharynx) for *O. nungara* (left) and *S. mediterranea* (right). Notably, the mucus fraction is clearly separated from the other samples in both species.

**Figure S4:** Barplot representing the summarized Blastp annotation results for the mucus fraction of *O. nungara* and *S. mediterranea*.

**Table S1**: Integrated results of all the putative toxins identified in *O. nungara* and *S. mediterranea*. Includes HOG information and blast summarized annotations for each putative toxin.

**Table S2**: Proteomics results for the toxins detected in *S. mediterranea*, including the normalized abundance of each sample.

**Table S3**: Proteomics results for the toxins detected in *O. nungara*, including the normalized abundance of each sample.

**Table S4:** Predicted toxins by Toxify and ToxinPred2 for the sequences included in HOGs that already contained at least one predicted and detected toxin in *S. mediterranea* or *O. nungara*.

**Table S5**: Pelican results and statistics.

**Table S6**: Integrated results of all the proteins identified in the mucus subset of *O. nungara* and *S. mediterranea*. Includes HOG information and blast summarized annotations for each mucus protein.

**Supplementary Materials** include all intermediate files, final files, scripts and tables, that can be found at the Github repository.

## Notes

https://github.com/MetazoaPhylogenomicsLab/Garcia_Vernet_et_al_2024_Flatworm_toxin_evolution

### Competing Interest Statement

The authors have declared no competing interest.

